# The AOPOntology: A Semantic Artificial Intelligence Tool for Predictive Toxicology

**DOI:** 10.1101/276832

**Authors:** Lyle D. Burgoon

**Author notes:** **Correspondence Address:** Lyle D. Burgoon, 3909 Halls Ferry Rd, Vicksburg, MS 39180.

## Abstract

**Introduction:** Toxicology needs artificial intelligence tools that can automate the prediction of toxicity. Today we are at an interesting nexus. We have thousands of chemicals in the environment that lack regulatory thresholds for determining risk. New high throughput in vitro testing methods are becoming available to test these chemicals. Causal Adverse Outcome Pathway Networks (CAOPN) are emerging that will allow us to make predictions based on perturbations of specific key events within the network. The AOPOntology was developed as infrastructure for this nexus, providing the ability to model and marry the data from the in vitro tests for the thousands of chemicals and place them within the CAOPN framework to facilitate adverse outcome predictions.

**Materials and Methods:** The AOPN is a functional specialized ontology that creates classes that model biological pathways and CAOPNs. Adverse outcome predictions are based on mathematical determinations of key events that are sufficient to infer adverse outcomes will occur, or biological information. These sufficiency relationships are captured in the AOPOntology and used by the semantic reasoners to make predictions.

**Results:** The AOPOntology version 1.0 architecture is in place, and a CAOPN for steatosis demonstrates how causal network theory is used to make predictions. The AOPOntology is available at https://github.com/DataSciBurgoon/aop-ontology.

**Discussion:** The AOPOntology is a knowledge base for CAOPNs that one can use to make predictions about a chemical’s potential toxicity using in vitro high throughput and other assays.

**Conclusions:** Using CAOPNs and causal network theory one is able to predict potential toxicity for chemicals using in vitro high throughput and various high content screens.

## Introduction

Every day the US military designs or acquires new chemicals, or faces challenges with existing legacy chemicals on its fields, ranges, and installations. For new chemicals, the challenge is to create new materials that are less toxic to humans and the environment. For legacy chemicals, the challenge is that safe exposure levels may not be known due to a lack of data.

Coupling in vitro high throughput screening assays with Adverse Outcome Pathways (AOPs) hold the promise of being a more ethical and relevant testing approach for identifying potentially toxic chemicals. Adverse Outcome Pathways are causal pathways that show the cascade from the point where a chemical interacts with a receptor (molecular initiating event) through the key events that are necessary for resulting in an adverse outcome at the individual and/or population level (1). There are three key stipulations to an AOP: 1) it must be linear, and 2) the key events within an AOP must be measurable, and 3) the key events must be high level events, such that many assays may map to a single key event. However, it has been our experience that in order to perform predictive toxicology using high throughput screening and high content assays, we need to combine several AOPs together into networks, and we need additional levels of granularity at the key event level.

Thus, I developed the concept of the Causal Adverse Outcome Pathway Networks (CAOPNs). These are more granular AOP networks that are more similar to molecular biological disease networks, such as those found in the Kyoto Encyclopedia of Genes and Genomes (KEGG), WikiPathway, and Reactome. Due to their causal network nature, one can generally apply causal network theory, specifically the backdoor analysis (2), to identify nodes/key events that are sufficient to infer an adverse outcome. The first step of the backdoor analysis algorithm is to introduce an edge between all parent nodes that share a child node. Next, the algorithm starts at the adverse outcome and identifies the shortest path to the molecular initiating event (MIE) of interest (note that in CAOPNs there may be more than one MIE). The first causal node found is the one just prior to the adverse outcome in the shortest path. Next, the algorithm removes the causal node from the pathway, and repeats this shortest path calculation, and identifies any other causal nodes. These steps are repeated until there are no more paths leading to the adverse outcome from the MIE.

In this work, I was interested in developing an artificial intelligence approach that could take in data about chemicals for specific CAOPNs and then predict potential adverse outcomes. This type of artificial intelligence is termed an expert system – it models expert knowledge about how chemicals cause adverse outcomes (the CAOPNs). One can encode what information we have about chemicals into the AOPOntology, and then use computational reasoners to apply first order logic across the data to make inferences about whether a chemical causes particular adverse outcomes. For this to work, one has to identify the key event(s) that are sufficient to infer the adverse outcome.

The AOPOntology is available for download from Github at https://github.com/DataSciBurgoon/aop-ontology.

## Materials and Methods

The AOPOntology builds on, or imports, several existing ontologies including ChEBI (3), human phenotype ontology (4), BioAssay Ontology (5), the uPheno ontology, and all of their associated ontologies. AOPOntology also includes linkages to the UniProt database to link proteins from the AOPOntology with UniProt. The AOPOntology is an OWL (web ontology language) ontology built using Protégé 5.

The design goal of the AOPOntology is to facilitate knowledge-based inference of potential adverse outcomes using chemical screening and prioritization data from high content and high throughput assays. I started with the AOP Framework as it was being implemented in the AOP-Wiki, and expanded it to include the ability to model and incorporate toxicological data from the high content and high throughput assays. Wherever possible, based on the literature, I defined sufficient key events – that is, those key events whose perturbation is sufficient to infer an adverse outcome. In addition, when that information was not available, the backdoor algorithm has been used to define those.

I have defined a small number of AOPs in the ontology based on CAOPN development in support of our military program. I have taken some AOPs from the AOP-Wiki, as well as others from the AOPXplorer (AOP-Wiki and AOPXplorer are both part of the international AOP-KB project, coordinated by the Organisation for Economic Cooperation and Development). I am developing software to automate the process of taking AOPs from the AOP-Wiki and AOPXplorer and encoding them into the AOPOntology.

## Results

The AOPOntology is available from Github at https://github.com/DataSciBurgoon/aop-ontology. The file aopo.owl contains the actual ontology. The other .owl files in the directory are used by the AOPOntology and required to open the aopo.owl file in Protégé 5.

The AOPOntology models Adverse Outcome Pathways (AOPs) as a parent class that contains classes for several different child AOP groupings. Currently, the AOPOntology contains the DevelopmentalToxicologyAOP, the DiseaseAOP, the LiverToxicityAOP, and the ReproductiveToxicologyAOP. Each of these classes have different AOPs associated with them, such as the NeuralTubeDefect AOP (under DevelopmentalToxicologyAOP) and DiabetesMellitusType2 (under DiseaseAOP).

From there, I developed specific instances of the different types of AOPs, for example, “AOP Neural Tube Defect via Hoxb1” is a specific instance of the NeuralTubeDefect AOP.

Members of the AdverseOutcomePathway class share characteristics. These include the fact that they must have exactly 1 adverse outcome, exactly 1 chemical molecular initiating event relationship (CMIER), and they can have any number of various types of AOP relationships (Figure 1). AOP relationships include the CMIE relationship and key event relationships (KERs). Figure 2 shows an example of how the linear components of the aromatase inhibition leading to loss of fecundity AOP are modeled by the AOPOntology.

**Figure 1:**
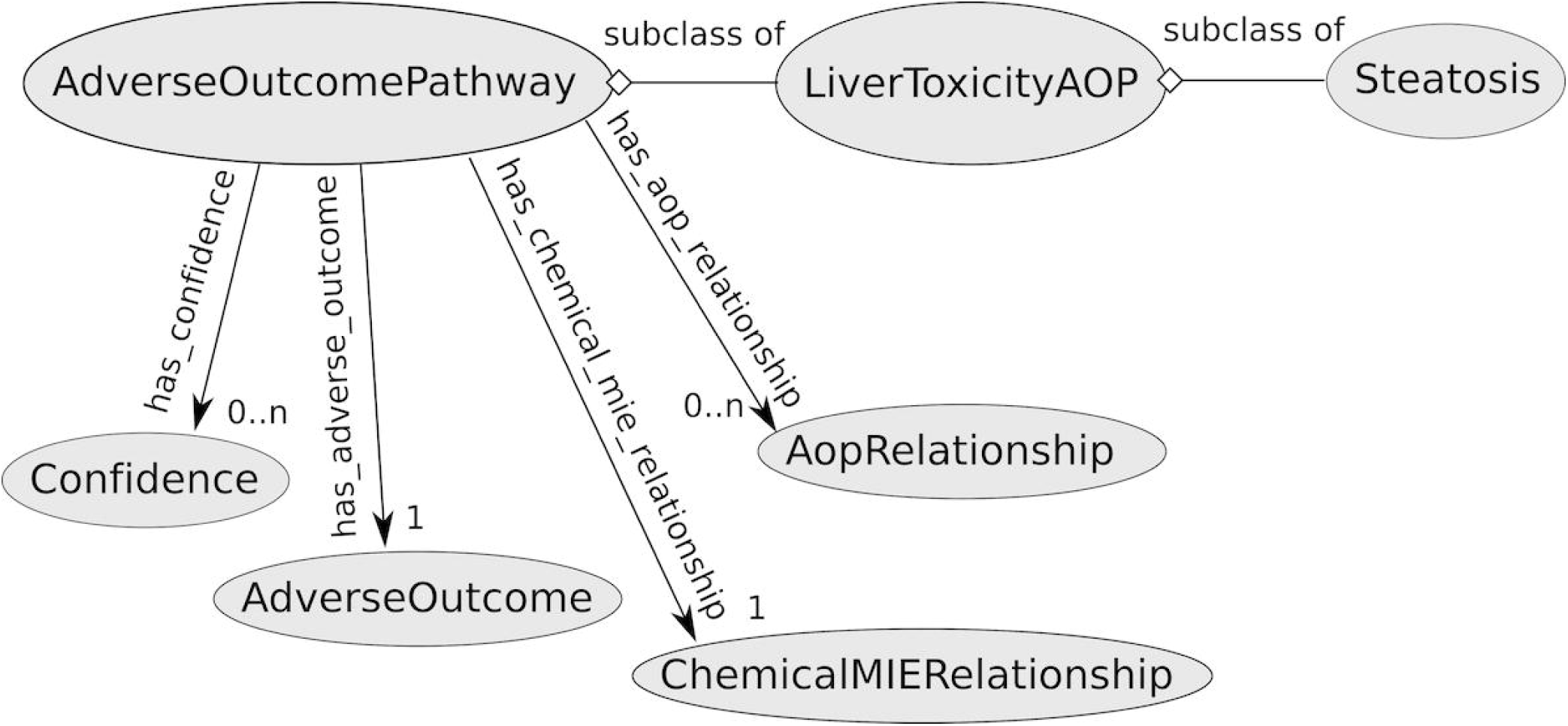
Overview of Adverse Outcome Pathway modeling in the AOPOntology. The AOPOntology models an Adverse Outcome Pathway as a collection of AopRelationships, where an AopRelationship defines a key event relationship (i.e., a relationship between upstream and downstream key events). Each AdverseOutcomePathway object also needs to have only one ChemicalMIERelationship – this is the relationship between a chemical and a molecular initiating event. The chemical can be left as an unknown, but it is essential that the ChemicalMIERelationship exists to demarcate the molecular initiating event (MIE). Each AdverseOutcomePathway object also must have only 1 AdverseOutcome. The Confidence in an Adverse Outcome Pathway is optional (denoted by 0‥n).

**Figure 2:**
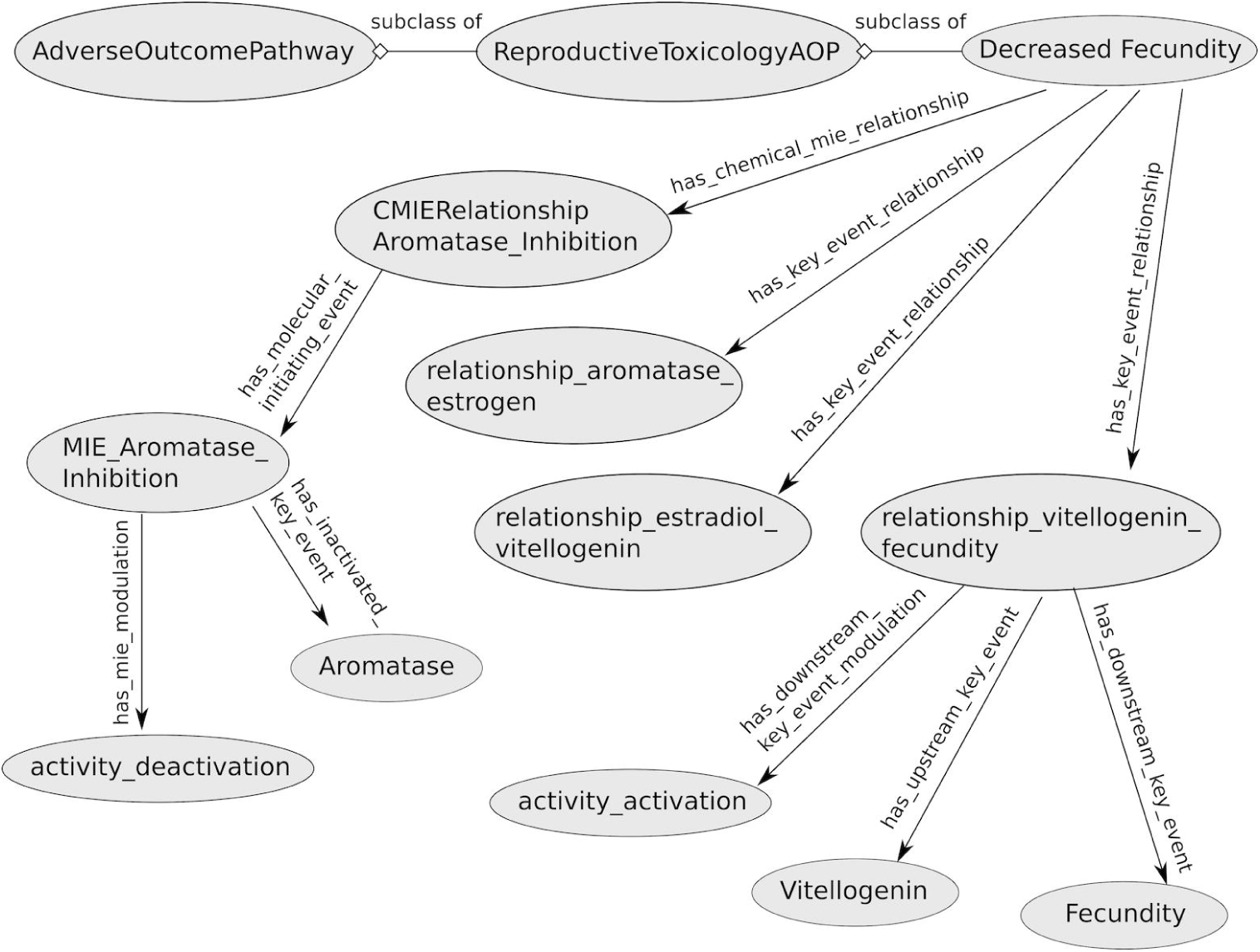
Example of how Key Event Relationships are modeled in the Inhibition of Aromatase Leads to Decreased Fecundity AOP. Here we can see that the Decreased_Fecundity AOP is a subclass of the ReproductiveToxicologyAOP, which is itself a subclass of AdverseOutcomePathway. Here we are only showing the AOPRelationships, which include the ChemicalMIERelationship, and several KeyEventRelationships (note that ChemicalMIERelationship and KeyEventRelationships are both subclasses of AopRelationship). This demonstrates how the linear AOP itself is captured in the AOPOntology.

Bioassay data can also be modeled within the AOPOntology. I have extended the BioAssay Ontology (BAO) to help us accomplish this. An example will make this clearer. I created an instance of oxidoreductase activity assay (from BAO) and called it hsd17b4_bioassay. Since I am using data from PubChem Assay ID 893, I have annotated hsd17b4_bioassay as having a PubChemAID of 893. Our instance of hsd17b4_bioassay also has object properties of “has participant” HSD17B4, “has participant” BkF (the chemical of interest in this example), “has measure group” and “has endpoint”. Since it has a participant of HSD17B4 this allows us to connect the assay directly to a key event within any of our CAOPNs.

In addition, the AOPOntology has assay calls. The has_chem_bio_assay_call object property has a ChemBioAssayCall, which can be one of either ActivatedAssayCall, InactivatedAssayCall, or NoChangeAssayCall. In our example, BkF results in an inactivated assay call. That inactivated assay call instance, BkF15umBayesNSMRCall has an object property of has_inactivated_key_event HSD17B4.

One can begin to make predictions using the assay calls. For this to work, the AOPOntology must have sufficiency arguments for as many of our CAOPNs as possible. When there is a lack of biological information, then one should use the backdoor algorithm to make an educated guess based on causal network theory. In this case, I have defined inactivation of HSD17B4 activity as sufficient to result in steatosis (6). The reasoners and query engines that use OWL are smart enough to understand that any time an instance contains “has_inactivated_key_event some HSD17B4” – in other words, any time one has a chemical that inactivates HSD17B4 activity according to one of our assays – then that chemical will lead to steatosis. Thus, we would predict that this chemical may cause steatosis (assuming ADME properties allow the liver to be exposed to the chemical).

## Discussion

The AOPOntology is a functional ontology geared at making computational toxicology predictions for hazard identification. Unlike other ontologies that the reader may be familiar with, the AOPOntology is not intended to be an encyclopedic reference. The AOPOntology is solely intended to make it possible to integrate high content and in vitro high throughput assays with CAOPNs to perform hazard identification.

One of the advantages to using the AOPOntology is that it will allow us to develop mathematically defensible and biologically sound assay batteries that minimize the total number of assays required to test for a full suite of adverse outcomes. By focusing on only those assays that are mathematically or biologically sufficient to infer adversity, one can decrease the overall testing burden while maximizing information content. This will lead to a net economic boost (through decreased testing requirements) and a net increase in chemical testing efficiency.

This paper briefly introduced some of the concepts that underlie the AOPOntology, and demonstrate its utility. Moving forward, I intend to populate the AOPOntology with the contents from the AOPXplorer, the AOP-KB, as well as data sources such as the Integrated Chemical Environment (ICE; https://ntp.niehs.nih.gov/pubhealth/evalatm/resources-for-test-method-developers/ice/), ToxCast, and PubChem.

Future work will include the development of graphical user interfaces to facilitate end-user querying of our systems, as well as the ability for users to upload their own data into the system. I see these as necessary next steps to make the AOPOntology useful to toxicologists.

In addition, I am using lessons learned from developing the AOPOntology to help develop foundational ontologies for developmental toxicology and zebrafish toxicity. These foundational ontologies will be more focused on being the encyclopedic ontologies that biologists are more familiar with, and will be less focused on artificial intelligence applications.

## Conclusions

The AOPOntology facilitates the development of minimal assay batteries and connecting in vitro high throughput screening and high content assay data to CAOPNs. This allows toxicologists to more quickly make mathematically defensible and biologically plausible hazard identifications in a more objective fashion.

## Acknowledgements

The author would like to acknowledge the work of Kyle Painter on early iterations of the Mode of Action Ontology, the predecessor to the AOPOntology.

## Author Disclosure Statement

The author has no conflicts of interest connected to this work.

## Correspondence Address

Lyle D. Burgoon 3909 Halls Ferry Rd Vicksburg, MS 39180

